# REDIRECTION: Generating drug repurposing hypotheses using link prediction with DISNET data

**DOI:** 10.1101/2022.07.26.501105

**Authors:** Adrián Ayuso Muñoz, Esther Ugarte Carro, Lucía Prieto Santamaría, Belén Otero Carrasco, Ernestina Menasalvas Ruiz, Yuliana Pérez Gallardo, Alejandro Rodríguez-González

## Abstract

In the recent years and due to COVID-19 pandemic, drug repurposing or repositioning has been placed in the spotlight. Giving new therapeutic uses to already existing drugs, this discipline allows to streamline the drug discovery process, reducing the costs and risks inherent to *de novo* development. Computational approaches have gained momentum, and emerging techniques from the machine learning domain have proved themselves as highly exploitable means for repurposing prediction. Against this backdrop, one can find that biomedical data can be represented in terms of graphs, which allow depicting in a very expressive manner the underlying structure of the information. Combining these graph data structures with deep learning models enhances the prediction of new links, such as potential disease-drug connections. In this paper, we present a new model named REDIRECTION, which aim is to predict new disease-drug links in the context of drug repurposing. It has been trained with a part of the DISNET biomedical graph, formed by diseases, symptoms, drugs, and their relationships. The reserved testing graph for the evaluation has yielded to an AUROC of 0.93 and an AUPRC of 0.90. We have performed a secondary validation of REDIRECTION using RepoDB data as the testing set, which has led to an AUROC of 0.87 and a AUPRC of 0.83. In the light of these results, we believe that REDIRECTION can be a meaningful and promising tool to generate drug repurposing hypotheses.

## I. Introduction

The increasing availability of tremendously large volumes of biomedical data obtained from improved multi-omics techniques has allowed science to redefine the way we conceive diseases, becoming more holistic and interrelated entities [1]– [3]. And more importantly, it has opened up new horizons in the possible treatments that can be used to heal the variety of pathologies categorized so far [4]. One of the terms that has been receiving rising attention, and especially considering the pandemic times we have lived in [5], is “drug repurposing or repositioning” [6]. The term refers to providing novel therapeutic uses to already existing drugs, that is, using them to treat other diseases different from the ones they were developed for [7].

Among the different approaches that have been suggested to tackle drug repurposing from the computational point of view [8], [9], one of the domains that could have a greater impact is Graph Machine Learning (GML), and specifically Graph Deep Learning (GDL) [10]. GDL or “deep learning on graphs” [11], [12] is a blooming field that, among its immense amount of applications, can indeed greatly contribute to link prediction tasks [13]. From the biomedical standpoint, predicting links in these biomedical networks can give invaluable insights. Particularly, potential connections between drugs and diseases not yet considered can be inferred [14], which unfolds a whole new world of more sophisticated paths regarding disease treatment and drug repurposing.

In the current work, we present a new model based on GDL called REDIRECTION (“dRug rEpurposing Disnet lInk pREdiCTION”). It has been trained with a part of the biomedical graph of the DISNET project (https://disnet.ctb.upm.es/) [15], consisting of diseases, symptoms, drugs, and the relationships between them. The main objective of REDIRECTION is to generate drug repurposing hypotheses by means of link prediction (also known as link inference). That is, the model’s output, once trained, provides a score ranging from 0 to 1 to every possible link in the graph, where 0 represents a null likelihood of that edge to exist in the graph, and 1, the total probability that the nodes are connected. The paper is organized as follows: Section II includes a brief revision of how GDL has been previously employed in the context of biomedical data, Section III details how the heterogeneous graph has been built, the specifications of REDIRECTION and how it has been evaluated and validated. Section IV depicts the different results obtained, while Section V finishes with the conclusions and future lines.

## II. Related work

The enormous volumes of biomedical data available nowadays have allowed science to study a wide range of interdisciplinary areas related to health [16], [17]. In particular, they have enhanced multiple phases of the drug discovery and development pipeline [18] reducing some of its disadvantages: high time-consumption, investment, and risks [19]. The interconnected nature of the biological information produced and utilized in drug discovery processes is a key property. The different interactions between the biological entities can be modelled as graphs representing relationships between drugs, side effects, diagnostic markers, treatments, test results, and so on. In this context, the goal of drug repurposing can be achieved either by identifying: (i) off targets with a potential action in the development of the disease due to drugs’ pleiotropic activity (off-target repurposing), (ii) similarities between diseases (on-target repurposing) or (iii) synergistic combinations of therapies (combination repurposing) [10]. Hereafter, we have exposed relevant GDL works on drug repositioning.

In off-target repositioning, some researchers employed protein structures to predict novel drug-target interactions. For instance, Torng and Altman proposed a GNN framework to predict drug-protein pockets associations. They represented drugs based on their chemical structures and protein pockets based on a set of key amino acid residues connected via Euclidean distance [20]. Moreover, multiple drug-target relationship inference works have been developed without using protein structures [21]–[24]. Olayan et al. introduced DDR to identify novel drug-target associations. First, they constructed a graph with drug and protein nodes connected by edges representing their similarity according to a heuristic from multiple data sources. DDR embedded each drug–protein pair based on the number of paths of predefined types that connected them within the graph. The resulting embeddings were fed to a random forest algorithm for drug–target prediction [23]. In 2020, Mohamed et al. presented an end-to-end knowledge graph embedding model to predict off-target interactions. The authors built a large knowledge graph enclosing drug and protein diverse data including pathways and diseases [24].

Semantic network completion approaches have been particularly effective in on-target repurposing tasks. Yang et al. introduced bounded nuclear norm regularization (BNNR), a block matrix and a disease-drug indication matrix. The method was based on the matrix completion property of singular value thresholding algorithm applied to the block matrix [25]. Alternatively, Wang et al. proposed a drug indication prediction method based on two bipartite graphs, capturing drug-target and disease-gene interactions as well as a protein-protein interaction (PPI) graph [26].

In diseases with complex aetiology or in which treatment resistance development is common, combination repurposing is especially effective. Decagon was the first combination therapy GML work. In it, Zitnik et al. used a multi-modal graph of drug– side effect–drug triplets and PPI interactions to model polypharmacy side-effects [27]. Achieving a similar level of accuracy, Deac et al. incorporated a knowledge graph and used a co-attention mechanism [28].

Further completing the list of active developments done during COVID-19 health crisis, it is important to highlight that multiple research groups have explored GML and GDL approaches to find drugs that could be repurposed for SARS-CoV-2 treatment [5], [29]–[31]. For instance, Zeng et al. used *RotatE* to identify repurposing hypotheses to treat SARS-CoV-2 from a large knowledge graph constructed from diverse data sources that included multiple associations between entities such as drugs, diseases, genes, and anatomies [30]. Moreover, in the field of rare diseases, Sosa et al. proposed a graph embedding method to detect repurposing candidates following a link prediction task [32].

## III. Materials and methods

We here present the process that we have followed to build the heterogeneous graph that served as input to train our model, REDIRECTION. We detail the model’s architecture, how it was trained, how we evaluated it, and how we performed a secondary validation with external standard data.

### A. Building the heterogeneous graph with DISNET data

The data employed to build the graph was obtained from the DISNET database (https://disnet.ctb.upm.es/), which is a biomedical integrated knowledge base containing information regarding diseases and their associations to symptoms and drugs, among others. In the case of disease-symptom relationships, DISNET gathers these data from processing unstructured textual sources (Wikipedia, PubMed and MayoClinic) by means of Natural Language Processing (NLP) [15]. The DISNET phenotypical information used for the present work was queried in September 2021. In the case of drugs and disease-drug relationships, DISNET information was previously integrated from the Comparative Toxicogenomics Database (CTD; http://ctdbase.org/), [33], [34] which is a robust service that aims to advance the understanding we have about diseases. From this source, DISNET uses the connections between diseases and chemicals specifically classified as “therapeutic”. The CTD integrated data in DISNET knowledge base was queried in May 2020.

The complete heterogeneous graph *G*(*V, E*) was formed as detailed hereunder. A schematic representation of its structure is included in Figure 1.

**Fig. 1.**
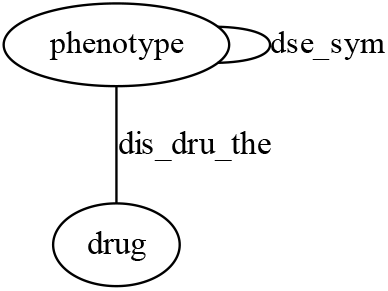
Structure of the heterogeneous graph.

- The set of nodes (*V*) consisted of two types: phenotypes (that is, diseases and symptoms) and drugs. There was a total of 34,673 nodes (30,729 phenotypes and 3,944 drugs). Phenotypes were codified by Unified Medical Language System (UMLS) [35], [36] Concept Unique Identifiers (CUIs), and drugs, by ChEMBL identifiers [37]– [39].
- The set of edges (*E*) consisted of 2 undirected types: ‘*dis_sym*’ links and ‘*dis_dru_the*’ links. On the one hand, the relationships between phenotypes, that is, between diseases and their symptoms, were named ‘*dis_sym*’ links. These edges could be regarded as intra-layer edges since they connected nodes in the same phenotypical layer. On the other hand, the relationships between diseases and the drugs that had a therapeutic effect on them were named ‘*dis_dru_the*’ links. These links could be considered as inter-layer edges since they connected nodes in the two different layers. There was a total of 313,972 ‘*dis_sym*’ links and 52,179 ‘*dis_dru_the*’ links.

These numbers varied when splitting the graph to train, evaluate and validate the model. It is worth noticing that in order to later perform the secondary validation of the developed model, we initially re oved so e ‘*dis_dru_the*’ edges from the graph before carrying out every partition. Those ‘*dis_dru_the*’ removed edges corresponded to the information present in RepoDB [40], which contains a standard set of drug repositioning successes and failures that can be used to benchmark computational repositioning methods. As explained in the next subsection, the connections between drugs and diseases that RepoDB offers were used to validate and interpret the results returned by the trained predictor. The idea was to, from drug repurposing cases that had already been tested or classified as successful, check whether the score provided by the model when inserting those links was higher than a randomly generated set of links.

### B. Predicting disease-drug links

#### I) REDIRECTION architecture

The model was developed under an encoder-decoder framework, meaning that an encoder firstly produced a vector embedding for each node in *G* and a decoder then performed a reconstruction of the information about each node with these embeddings to predict ‘*dis_dru_the*’ links. The model was based on the Graph Neural Networks (GNNs) [41] formalism and its architecture is depicted in Fig. 2.

**Fig. 2.**
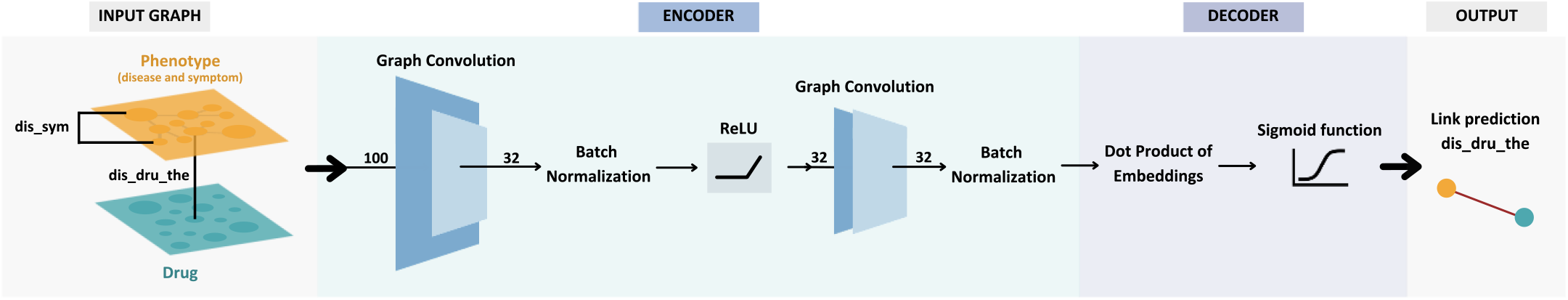
Representation of REDIRECTION model architecture.

For the encoder, we used a two-layer convolution approach. After the first layer, messages were passed through batch normalization and submitted to a non-linearity activation function (ReLU). Each layer has been based and shares parameters (except from the hidden dimensions) with an approximation of GraphSAGE layers to heterogeneous graphs [42]. GraphSAGE generates embeddings by sampling and aggregating features from a node’s local neighborhood and the node’s features. In our case, as initial node’s features for the two types of nodes, we used input feature vectors *x*_*v*_, so that

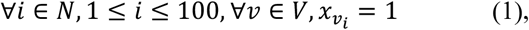

denoting that each *x*_*v*_ associated to each node in *G*, was formed by 100 dimensions with value 1. For the decoder, we employed the dot product, that, given two embeddings A and B, is computed as detailed hereunder:

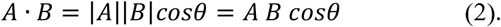

It allows reconstructing the relationships between the node embeddings previously generated by the encoder. The dot product returns coefficients, which are in turn passed to a sigmoid function to normalize them in the range [0,1]. In this manner, we can understand the output score as the weight that REDIRECTION gives to the links in the graph, where 1 represents the higher feasibility of the existence of the edge.

REDIRECTION has been developed in Python 3.8.10, under the framework of the Stanford library DeepSNAP 2.0 (https://snap.stanford.edu/deepsnap/), which in turn uses PyTorch Geometric 2.0.3 and NetworkX 2.6.3. We made use of CUDA Toolkit 11.3, running the experiments on an Ubuntu Server LTS 20.04.4 with a GPU (NVIDIA GeForce RTX 3090 24GB). All code and results have been published in an accessible repository^1^.

#### 2) Training

To train the model, the specific training set, formed by the 80% of the data of the original graph, was employed. From the remaining 20% of the data, a 10% was assigned for internal training validation tasks (i.e., to avoid overfitting, to tune the hyperparameters, and to ensure the absence of data leakage) and a 10% for testing tasks.

During this training phase, we optimized the model parameters by using the Binary Cross Entropy with Logit Loss (BCELogitLoss) as the loss function, which is computed as follows:

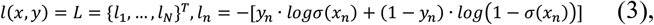

being *x*_*n*_ the model’s prediction and *y*_*n*_ the true label (referring to whether the edge exists or not). We performed a negative sampling [43], [44], that is, for each edge in the graph (positive edge), we sampled a random edge (negative edge) that was not present in the original graph.

We optimized the model with the Adam optimizer [45], through 200 epochs (training iterations) and a weight decay of 1e^-4^. We used a learning rate of 0.01 and a ratio for the edge message passing of 0.8. The dimension of the “hidden e beddings” (those generated after the first convolution) was 32. We approached the problem as an end-to-end optimization, so we optimized all the trainable parameters in tandem and propagated the loss function gradients through the encoder and the decoder.

#### 3) Evaluating and validating REDIRECTION

We envisioned the evaluation and validation of the developed predictor in two separated steps. We would like to pinpoint that there was not information leakage between the different subgraphs used for the training, the testing, and the secondary validation.

Firstly, we used the testing set, formed by a 10% of the original graph, to obtain classical evaluation metrics. Those are the Area Under the Receiver Operators Characteristics Curve (AUROC) and the Area Under the Precision-Recall Curve (AUPRC), as well as the visualization of both curves themselves. The first is based on the number of True Positive Rate (TPR) and False Positive Rate (FPR). The second one measures the tradeoff between precision and recall for different thresholds, and it is preferred when the dataset is imbalanced.

Secondly, we performed a further validation with real cases of drug repurposing. Those were obtained from RepoDB (more information on this source is included in the last paragraph of subsection III.A). The data coming from RepoDB that was already included in the DISNET knowledge base was removed from the original graph in order to fairly prevent any kind of data leakage between the training stage and this validation. A total of 5,013 disease-drug connections formerly tested or classified as successful in the context of repurposing were taken. This validation links followed the same structure as the edges provided for training, meaning that each of them was formed by a disease identified with its UMLS CUI and a drug identified with its ChEMBL ID.

Moreover, we also compared the distribution of the scores returned by REDIRECTION for RepoDB links against the ones predicted for a random set of the same number of links randomly picked. The idea behind this comparison was to observe whether our model was capable or not of assigning scores closer to 1 to those edges coming from real repurposing cases than to the rest of possible links in the graph.

## IV. Results and discussion

After the evaluation with the testing subset, REDIRECTION model showed an AUROC of 0.93 and a AUPRC of 0.90. When validating the model with RepoDB actual disease-drug links, it obtained an AUROC of 0.87 and a AUPRC of 0.83. Fig. 3 depicts the ROC and Precision-Recall curves for this experiment.

**Fig. 3.**
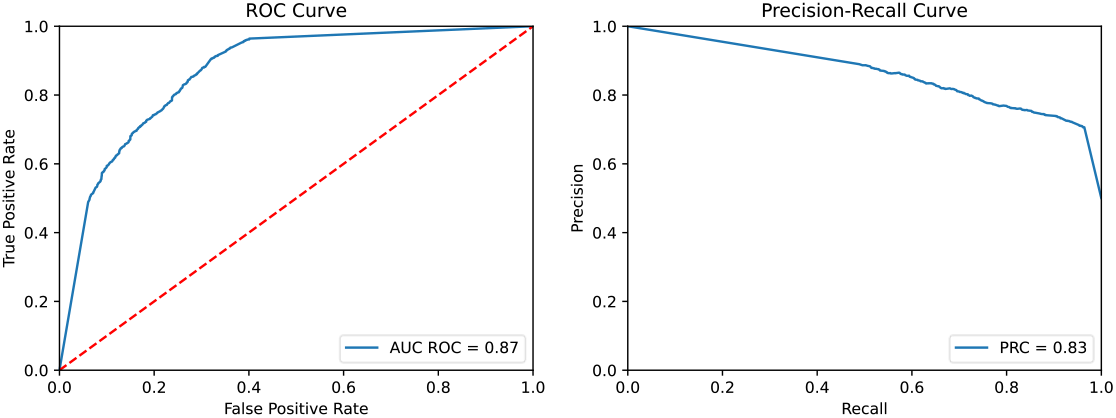
Receiver Operating Characteristics (left) and Precision-Recall (right) curves obtained from REDIRECTION when validated with RepoDB subset.

Therefore, the acuracy observed when putting REDIRECTION to the test can be regarded as more than reasonable considering that just a small part of the DISNET complete graph was used to train it (alluding to the fact that not all types of biomedical entities were included for the time being).

The scores returned when validating with RepoDB links presented an average value of 0.69 and a standard deviation of 0.44. Table I includes 20 of the RepoDB disease-drug links that have been scored with a 1 by REDIRECTION. We could interpret that these edges could have stronger evidence of underlying structures and connections in the DISNET subgraph. This means that if REDIRECTION has been able to predict such links with just the information of diseases, symptoms, drugs and their relationships, these entities could have a significant role to repurposing in their specific cases.

**TABLE I.** Top 20 examples of some of the links predicted with score=1 when validating REDIRECTION with RepoDB subset.

| Disease UMLS CUI | Disease Name | Drug ChEMBL ID | Drug name |
| --- | --- | --- | --- |
| C0030193 | Pain | CHEMBL1201193 | LEVOBUPIVACAINE |
| C0027819 | Neuroblastoma | CHEMBL1351 | CARBOPLATIN |
| C0036337 | Schizoaffective Disorder | CHEMBL111 | RIMONABANT |
| C0036341 | Schizophrenia | CHEMBL111 | RIMONABANT |
| C0011860 | Diabetes Mellitus, Non-Insulin-Dependent | CHEMBL111 | RIMONABANT |
| C0242339 | Dyslipidemias | CHEMBL111 | RIMONABANT |
| C0028754 | Obesity | CHEMBL111 | RIMONABANT |
| C0037299 | Skin Ulcer | CHEMBL1382627 | SULFADIAZINE, SILVER |
| C0033975 | Psychotic disorder | CHEMBL1621 | PALIPERIDONE |
| C1269683 | Depressive Disorder, Major | CHEMBL1201222 | LISDEXAMFETAMINE |
| C0024305 | Lymphoma, Non-Hodgkin | CHEMBL1351 | CARBOPLATIN |
| C1621958 | Glioblastoma Multiforme | CHEMBL1351 | CARBOPLATIN |
| C0007131 | Carcinoma, Non-Small-Cell Lung | CHEMBL1351 | CARBOPLATIN |
| C0035335 | Retinoblastoma | CHEMBL1351 | CARBOPLATIN |
| C1140680 | Malignant neoplasm of ovary | CHEMBL1351 | CARBOPLATIN |
| C0917801 | Insomnia | CHEMBL911 | ZOLPIDEM |
| C0011860 | Diabetes Mellitus, Non-Insulin-Dependent | CHEMBL385517 | SAXAGLIPTIN |
| C0018802 | Cardiac failure congestive | CHEMBL184412 | DRONEDARONE |
| C0004238 | Atrial Fibrillation | CHEMBL184412 | DRONEDARONE |

In the light of these results, it must be noted that not all the RepoDB edges were predicted by REDIRECTION as to be close to 1, but the majority was. And this makes us think that, even though the model behaves quite well, including other relevant biomedical entities to the training graph (as genes or proteins, for instance) could greatly contribute to enrich the graph structure. We hypothesize that this might lead REDIRECTION to predict even better over the RepoDB subset, and by extension, over other cases.

Finally, we include a comparison between these scores obtained with RepoDB and others obtained with a randomly generated set of links. We depict the distribution of both of them in Fig. 4. The density curve representing the distribution of the random edges’ scores (in blue) is clearly shifted towards 0, while the RepoDB score’s distribution presents a significantly higher number of links around value 1. Such a difference evidences the distinct nature of the REDIRECTION predictions and how it has been capable to distinguish between edges, giving higher scores to real repurposing cases.

**Fig. 4.**
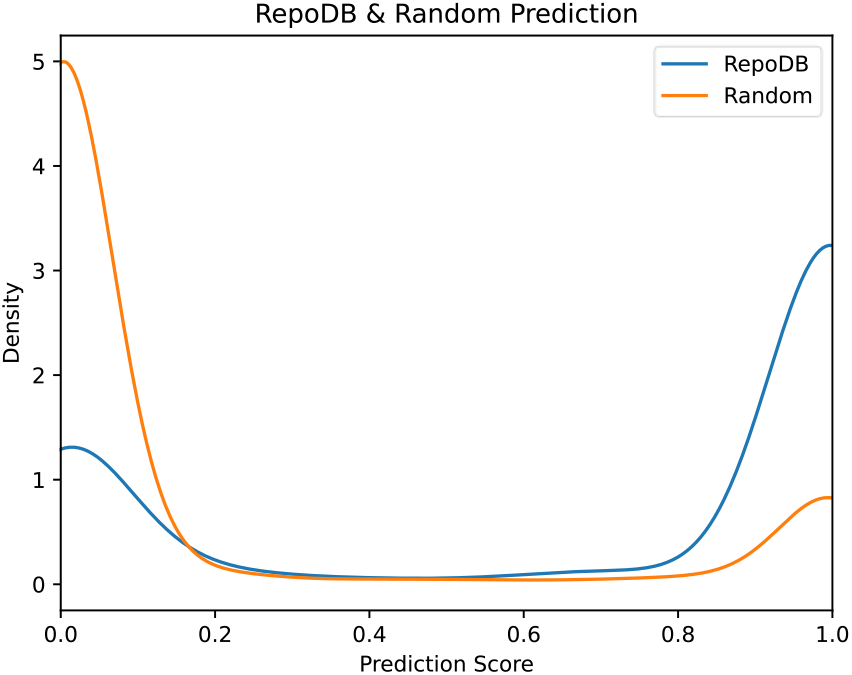
Comparison of the distribution of the score returned by REDIRECTION when having as input the RepoDB validating set (in blue) and a randomly generated set of links (in orange).

## V. Conclusions and future lines

The integration of heterogeneous biomedical information in the form of a multilayered, inter and intra-connected graph has become a fundamental approach to better understand diseases and their underlying elements and features. One of the most relevant disciplines in this computational context is drug repurposing, which tries to find new uses for already existing drugs. In the present work, we have presented a new model in the scope of DISNET biomedical graph, which aims to generate drug repurposing hypotheses. This model was called REDIRECTION and has been based on graph deep learning frameworks and link prediction tasks. To the best of our knowledge, the main advancement that we have introduced in the current research is that we have trained the predictor with a graph formed by diseases, symptoms, and drugs simultaneously. And it has performed fairly well when testing it both with the subset intended for it and with a real compilation of already-tested repurposing links from RepoDB.

As for the future lines, the most immediate avenue that this investigation opens is the development of more accurate and comprehensive models that are able to embed a larger heterogeneous biomedical graph. That is, the DISNET graph including other types of nodes not limited to phenotypes and drugs, such as genes, proteins, metabolic pathways, drug-drug interactions, and so on. It has to be mentioned that REDIRECTION is still a prototype and aims to be improved and refined by adding more information of different nature to the input graph, as well as by including other extended details to its architecture.

## Acknowledgment

The work is a result of the project “Data-driven drug repositioning applying graph neural networks (3DR-GNN)”, that is being developed under grant “PID2021-122659OB-I00” from the Spanish Ministerio de Ciencia, Innovación y Universidades. LPS’s work is supported by “Progra a de fomento de la investigación y la innovación (Doctorados Industriales)” fro Co unidad de Madrid (grant “IND2019/TIC-17159”). BOC’s work is supported by “For ación de Personal Investigador” grant “FPI PRE2019-090912” as part of the project “DISNET (Creation and analysis of disease networks for drug repurposing from heterogeneous data sources applied to rare diseases)”, developed under grant “RTI2018-094576-A-I00” from the Spanish Ministerio de Ciencia, Innovación y Universidades.

## Footnotes

1 https://medal.ctb.upm.es/internal/gitlab/disnet/cbms2022-gnns

